# Astrocytes amplify neuronal dendritic volume transmission

**DOI:** 10.1101/361014

**Authors:** Chun Chen, ZhiYing Jiang, Xin Fu, Jeffrey G. Tasker

## Abstract

In addition to their support role in neurotransmitter and ion buffering, astrocytes directly regulate neurotransmission at synapses via local bidirectional signaling with neurons. Here, we reveal a new form of neuronal-astrocytic signaling that transmits retrograde dendritic signals to upstream neurons to activate recurrent synaptic circuits. Norepinephrine activates α_1_-adrenoreceptors in hypothalamic corticotropin releasing factor (CRF) neurons to stimulate dendritic release, which triggers an astrocytic calcium response and release of ATP; ATP stimulates action potentials in upstream glutamate and GABA neurons to activate recurrent excitatory and inhibitory synaptic circuits to the CRF neurons. Thus, norepinephrine activates a novel retrograde signaling mechanism in CRF neurons that engages astrocytes in order to extend dendritic volume transmission to reach distal presynaptic glutamate and GABA neurons, thereby amplifying volume transmission mediated by dendritic release.

## Introduction

The functional role of astrocytes is not limited to buffering extracellular ions and neurotransmitters, but also includes an active involvement in neurotransmission. Astrocytes interact with neurons by responding to neurotransmitters and by releasing gliotransmitters ^1^. However, the neuron-astrocyte interaction has been largely limited to the tripartite synapse, or neuronal pre- and postsynaptic elements and surrounding astrocyte (Araque et al., 1999; Halassa et al., 2007). Astrocytic calcium signals can be transmitted through branched astrocyte arbors (Cornell-Bell et al., 1990; Charles et al., 1991; Porter and McCarthy, 1996) or can be localized to subdomains within branches (Haustein et al., 2014; Shigetomi et al., 2013), giving astrocytes the potential to signal remotely from the response generation site, and electrical coupling via gap junctions extends this capacity beyond the limits of the astrocytic arbor ^9^. Whether astrocytes transmit neuron-derived signals distally is currently not known.

Volume neurotransmission is mediated by diffusion of neurotransmitter away from its release site, and can be orthograde, between the presynaptic terminal and postsynaptic soma/dendrite, or retrograde, from the postsynaptic dendrite to the presynaptic neuron. It is spatially constrained by astrocytic buffering and enzymatic degradation (Di et al., 2013; Wu and Tasker, 2017; Son et al., 2013). Pioneering research on hypothalamic neuroendocrine cells demonstrated retrograde volume transmission of the neuropeptides oxytocin and vasopressin, which exerts localized paracrine actions on presynaptic neurotransmitter release (Ludwig et al., 2002; Kombian et al., 1997; Oliet et al., 2007; de Kock et al., 2003). We reported that vasopressin released from vasopressin neuron dendrites activates a calcium signal in astrocytes that leads to stimulation of presynaptic GABA neurons (Haam et al., 2014).

The hypothalamic-pituitary-adrenal (HPA) axis comprises the neuroendocrine stress response, producing systemic glucocorticoid secretion from the adrenals following corticotropin-releasing factor (CRF) secretion from the hypothalamic paraventricular nucleus (PVN) and adrenocorticotrophic hormone secretion from the pituitary. Brainstem noradrenergic systems provide a major excitatory drive to the HPA axis. Anatomical and physiological studies indicate that CRF neurons receive *direct* noradrenergic innervation (Itoi et al., 1994; Plotsky, 1987; Cole and Sawchenko, 2002; Itoi et al., 1999; Helmreich et al., 2001; Sawchenko and Swanson, 1982; Cunningham and Sawchenko, 1988; Flak et al., 2014; Woulfe et al., 1990; Flak et al., 2009; Herman et al., 2004; Ziegler et al., 2012), and medial parvocellular neurons express α_1_-adrenoreceptors (Cummings and Seybold, 1988; Day et al., 1997). In contrast, electrophysiological studies indicate that norepinephrine excites parvocellular neurons *indirectly* via activation of local synaptic circuits (Daftary et al., 2000; Han et al., 2002). Thus, despite multiple efforts, the mechanism of the noradrenergic activation of the HPA axis remains elusive.

Here, we investigated the cellular mechanisms of the norepinephrine excitation of CRF neurons using whole-cell recordings in brain slices. We found that norepinephrine elicits the dendritic release of vasopressin from the CRF neurons to activate an astrocytic relay to presynaptic glutamate and GABA neurons, revealing a previously unidentified astrocytic amplification of retrograde volume transmission.

## Results

### NE activation of CRF neurons

Extracellular loose-seal, cell-attached patch clamp recordings were performed in CRF neurons to record the spontaneous spiking activity. Bath application of norepinephrine (NE, 100 μM, 5 min) significantly increased the firing frequency of CRF-eGFP neurons (p < 0.01), which was suppressed in the glutamate receptor antagonists DNQX (15 μM) and AP5 (50 μM) (p = 0.37) and enhanced in picrotoxin (50 mM), though this did not reach significance (Two-way ANOVA, F (2, 33) = 3.522, p = 0.04, followed by Bonferroni’s multiple comparisons) (Fig. 1A-C). To test whether the excitation of CRF neurons is mediated by NE modulation of synaptic inputs, we recorded spontaneous EPSCs (sEPSCs) and IPSCs (sIPSCs) in CRF neurons. sEPSCs had a basal mean frequency of 1.56 ± 0.10 Hz, amplitude of 19.13 ± 0.50 pA, and decay time of 2.18 ± 0.06 ms (n = 47); bath application of NE (100 μM, 5 min) evoked a robust increase in the frequency of sEPSCs (335.2 ± 61.5% of baseline at 100 μM, n = 17, p < 0.01), without causing a change in sEPSC amplitude (p = 0.14) or decay time (p = 0.58) (Fig. 1D, E; Supplemental Fig. 2A), which suggested a possible presynaptic site of action. The NE stimulation of excitatory synaptic inputs to CRF neurons was concentration-dependent, with a threshold concentration of ~100 nM (118.7 ± 7.99% of baseline, n = 9, p = 0.047) (Fig. 1F). It was also action potential-dependent, since blockade of voltage-gated Na^+^ channels with tetrodotoxin (TTX, 1 μM) abolished the NE-induced increase in sEPSC frequency (92.4 ± 9.0% of baseline, n = 9, p = 0.42) (Fig. 1G).

**Figure 1.**
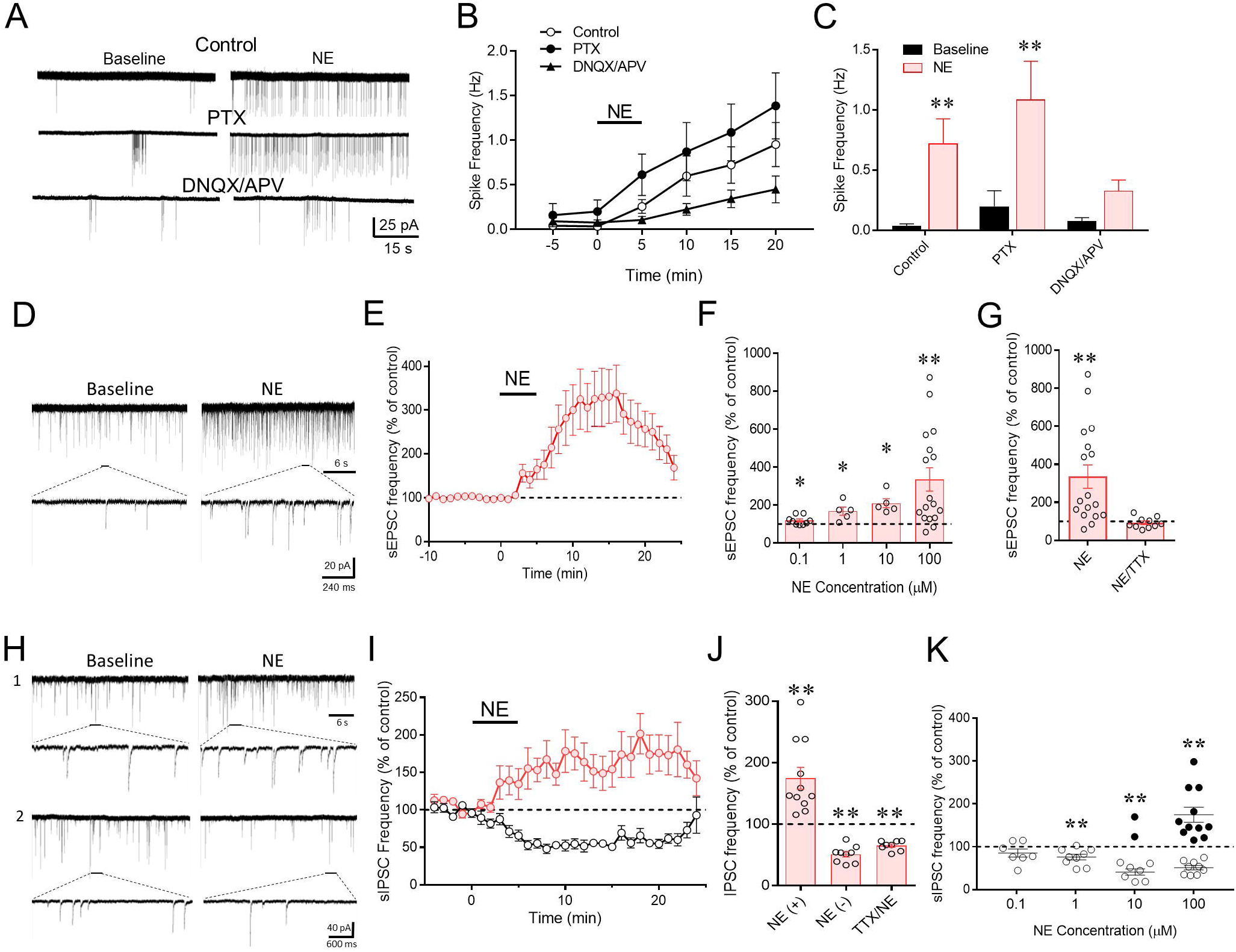
Norepinephrine regulation of CRF neurons via local synaptic circuits. A. Loose-seal patch clamp recording of the spiking response of a CRF neuron to bath application of NE (100 μM) and the increased response following blockade of GABA_A_ receptors with picrotoxin (PTX) and decreased response following blockade of glutamate receptors with DNQX/APV. B. Time series of changes in spike frequency prior to (−5 – 0 min), during (0 – 5 min), and after NE application (100 μM) in control solution, PTX, and DNQX/APV. C. Mean spike frequency responses to NE without (Control) and with the GABA_A_ (PTX) and glutamate receptor antagonists (DNQX/APV). D. Whole-cell recording of the NE (100 μM) effect on sEPSCs in a CRF neuron recorded in bicuculline to block inhibitory synaptic currents. E. Time plot of the NE (100 μM) effect on the frequency of sEPSCs. F. The NE effect was concentration-dependent, with a threshold concentration ~ 0.1 μM. G. The NE-induced increase in sEPSC frequency was blocked by TTX, suggesting spike-dependence. H. Whole-cell recording of the NE (100 μM) effect on sIPSCs in a CRF neuron recorded in DNQX/APV to block excitatory synaptic currents. Norepinephrine caused an increase in sIPSCs in some cells (1) and a decrease in sIPSCs in others (2). I. Time plots of the normalized changes in mean sIPSC frequency in response to NE in different cohorts of CRH neurons. J. Norepinephrine either increased or decreased the frequency of sIPSCs relative to baseline, suggesting presynaptic sites of modulation. The NE-induced increase, but not the decrease, in sIPSC frequency was blocked by TTX, suggesting the facilitation, but not the suppression, of sIPSCs by NE was spike-dependent. K. The two sIPSC responses to NE had differing concentration dependencies, with the NE-induced suppression of sIPSCs dominant at lower concentrations and the NE-induced facilitation emerging in some, but not all recorded cells at higher concentrations of NE.

The sIPSCs in CRF-eGFP neurons had a basal mean frequency of 0.86 ±0.10 Hz, amplitude of 51.23±3.82 pA, and decay time of 12.72 ±0.54 ms (n = 20). Norepinephrine (100 μM, 5 min) induced two distinct effects on sIPSCs in separate cohorts of CRF neurons: 55% of CRF neurons (11/20) showed a ~75% increase in sIPSC frequency (p < 0.01) and 45% (9/20) showed a ~50% decrease in sIPSC frequency (p < 0.01) (Fig. 1H, I), with no effect on sIPSC amplitude (p = 0.53) or decay time (p = 0.12) (Supplementary Fig. 2B). The NE-induced increase in sIPSC frequency was not seen in any CRF neurons recorded in the presence of TTX, and all of the cells showed a decrease in the sIPSC frequency (65.14% ± 2.78%, n = 8, p < 0.01) (Fig. 1J). Thus, the NE facilitation of inhibitory synaptic inputs to CRF neurons is action potential-dependent, while the NE-induced suppression of inhibitory synaptic inputs is action potential-independent. The facilitatory effect of 100-μM NE on GABA release in some cells, therefore, masked the NE-induced suppression of GABA release, which occurred in all CRF neurons tested. The absence of the spike-dependent facilitatory response in some CRF neurons suggested that the axons of the NE-sensitive presynaptic neurons were cut in those slices.

The two effects of NE on synaptic inhibition had different concentration sensitivities. Nearly all the CRF neurons (14/16 cells) responded to NE with either no response or a suppression of sIPSCs at the lowest concentrations tested (100 nM and 1 μM), and the proportion of cells that responded with a facilitation of sIPSCs increased at higher NE concentrations (2/9 cells (22%) at 10 μM and 11/20 cells (55%) at 100 μM) (Fig. 1K). Thus, the spike-dependent NE facilitation of GABA release had a higher threshold (~10 μM) than the spike-independent NE suppression of GABA release (< 100 nM). Note that the threshold for facilitation of excitatory synaptic inputs (~100 nM) was over an order of magnitude lower than the threshold for NE facilitation of inhibitory synaptic inputs to the CRF neurons. Additionally, the NE activation of inhibitory synaptic inputs was less robust than the activation of excitatory synaptic inputs, which accounts for the net excitatory effect of NE on the CRF neuron spiking.

### α_1_-adrenoreceptor dependence of the norepinephrine facilitation of excitatory synaptic inputs to CRF neurons

We next investigated the adrenoreceptor dependence of the NE facilitation of excitatory synaptic inputs to CRF neurons. Bath application of the α_1_-adrenoreceptor antagonist prazosin (10 µM) alone caused a small decrease in sEPSC frequency (85.2 ± 5.5% of baseline, n = 9, p = 0.027), suggesting a tonic activation of α_1_ adrenoreceptors by ambient endogenous NE (or epinephrine). Prazosin abolished the NE-induced increase in sEPSC frequency (p = 0.35, n = 9) (Fig. 2A, C). Bath application of the α_1_ adrenoreceptor agonist phenylephrine (100 µM) for 5 min caused an increase in sEPSC frequency (424.2 ± 106.2% of baseline, n=8, p=0.019) similar to that induced by NE (NE vs. PE, p=0.45), although it did not reverse with a 20-min washout (Fig. 2B, C). Taken together, these results indicate that the NE-induced facilitation of sEPSCs in CRF neurons is mediated by α_1_-adrenoreceptor activation.

**Figure 2.**
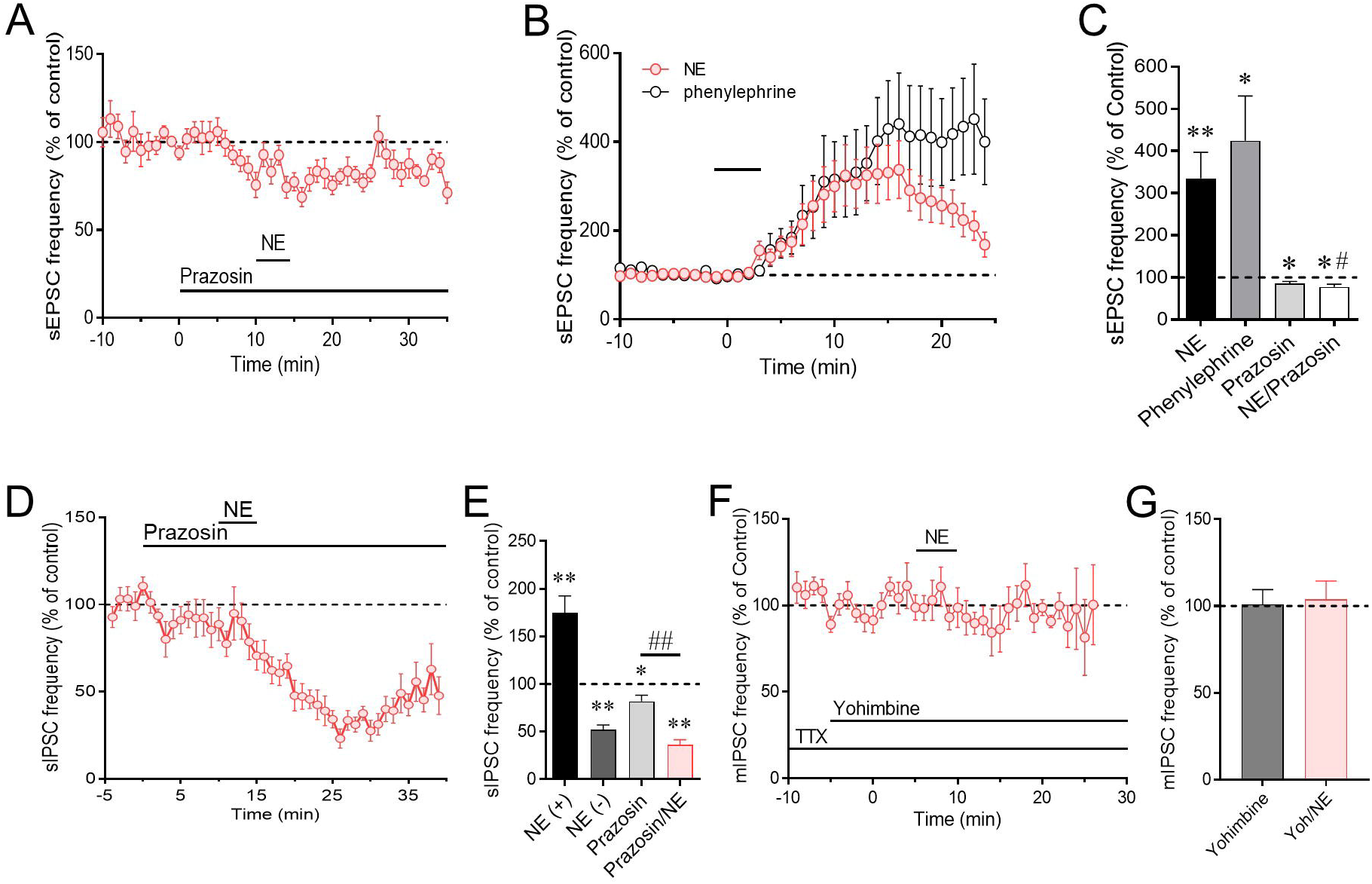
Adrenoreceptor dependence of norepinephrine effects on excitatory and inhibitory synaptic inputs to CRF neurons. A. The NE-induced facilitation of sEPSCs was blocked by the α1 receptor antagonist prazosin. B. The α1 receptor agonist phenylephrine, like norepinephrine, increased sEPSC frequency. C. Summary of α1 receptor agonist and antagonist effects on sEPSC frequency. D. The α1 receptor antagonist prazosin blocked the norepinephrine-induced increase in sIPSC frequency, but not the decrease in sIPSC frequency. E. Summary of α1 receptor antagonist effect on the facilitation and suppression of sIPSCs. F. The α2 receptor antagonist yohimbine blocked the norepinephrine-induced decrease in sIPSC frequency isolated in TTX. G. Summary of α2 receptor antagonist effect on NE-induced suppression of sIPSCs.

### α_1_- and α_2_-adrenoreceptor dependence of the norepinephrine modulation of inhibitory synaptic inputs to CRF neurons

In recordings of sIPSCs in the presence of DNQX and AP5, bath application of the α_1_ adrenoreceptor antagonist prazosin (10 μM) alone caused a significant decrease in the sIPSC frequency compared to baseline (81.82% ± 6.33% of baseline, n = 8, p = 0.02), again suggesting a tonic activation of α_1_-adrenoceptors and modulation of GABA release by endogenous NE. Norepinephrine (100 μM) failed to elicit an increase in sIPSC frequency following preapplication of prazosin, but all 8 CRF neurons recorded in prazosin responded with a significant decrease in sIPSC frequency (36.31% ± 5.01% of baseline, n = 8, p < 0.01) (Fig. 2D, E). This indicated that the NE facilitation of IPSCs, like the facilitation of EPSCs, is mediated by α_1_-adrenoreceptor activation, whereas the NE-induced suppression of IPSCs is α1-receptor independent. The NE-induced suppression of sIPSCs was greater in the presence of prazosin (51% of baseline without prazosin vs. 36% of baseline with prazosin) and this difference showed a strong trend towards significance (p=0.050) (Fig. 2E), which supports a model of opposing NE regulation of GABA release onto CRF neurons.

Because the NE facilitation, but not suppression, of GABA release was spike-dependent, we blocked the facilitatory response with TTX (1 μM) to isolate the NE-induced suppression of GABA release. In the presence of TTX, preapplication of the α_2_ adrenoreceptor antagonist yohimbine (20 μM) alone had no effect on sIPSC frequency, but completely blocked the NE-induced decrease in sIPSC frequency (103.8% ± 10.57%, n = 8, p = 0.86) (Fig. 2F, G). Thus, at lower concentrations, NE activates presynaptic α_2_ adrenoreceptors to suppress GABA release and, at higher concentrations, it also activates α1 adrenoreceptors to stimulate spiking in presynaptic GABA neurons and an increase in GABA release onto PVN CRF neurons.

### Localization of adrenoreceptors to pre- and postsynaptic loci

Our data suggest that NE causes a spike-dependent increase in excitatory and inhibitory synaptic inputs to PVN CRF neurons by acting at α1 adrenoreceptors on local presynaptic glutamate and GABA neurons, respectively. However, considerable immunohistochemical evidence exists for noradrenergic synapses directly on CRF neurons (Flak et al., 2009; Liposits et al., 1986) and for α_1_-adrenoceptor expression by CRF neurons ^34^. Here, we tested for the dependence of the NE effect on the activation of postsynaptic receptors by including a broad-spectrum G-protein inhibitor, GDP-β-S (1 mM), in the patch electrode. The NE-induced increase in sEPSC frequency was blocked in 7 of 10 CRF neurons (70%) recorded with GDP-β-S-containing electrodes (123.4 ± 36.6% of baseline, n = 10, p = 0.54) (Fig. 3A, B), which suggested that the α1 adrenoceptor-induced facilitation of excitatory synaptic inputs to the CRF neurons has a postsynaptic locus. With GDP-β-S in the patch solution and glutamate receptors blocked, only 15% of CRF neurons (2/13) responded to NE (100 μM) with an increase in sIPSC frequency, while 85% of CRF neurons (11/13) responded with a decrease in sIPSC frequency (71.53% ± 10.92% of baseline, n = 13, p = 0.023) (Fig. 3C, D). The shift in the distribution of the two sIPSC responses caused by postsynaptic G-protein blockade (increase: 55% without GDP-βs to 15% with GDP-βs; decrease: 45% without GDP-βs to 85% with GDP-βs) was significant (p = 0.023, chi-squared). This suggested that the NE-induced facilitation of GABA release, like that of glutamate release, is dependent on postsynaptic G protein activation, whereas the NE-induced suppression of GABA release is not. Therefore, the NE facilitation of excitatory and inhibitory synaptic inputs to CRF neurons share a common mechanism: postsynaptic α_1_ receptor activation that results in spike generation in presynaptic glutamate and GABA neurons. The NE-induced suppression of inhibitory synaptic inputs to the CRF neurons, on the other hand, is mediated by the activation of presynaptic α_2_ adrenoreceptors and is spike-independent, indicating that it occurs at presynaptic GABA terminals. The reliance of the NE facilitation of glutamate and GABA inputs on a postsynaptic G protein-dependent mechanism suggests the recruitment of a retrograde signaling mechanism in this effect.

**Figure 3.**
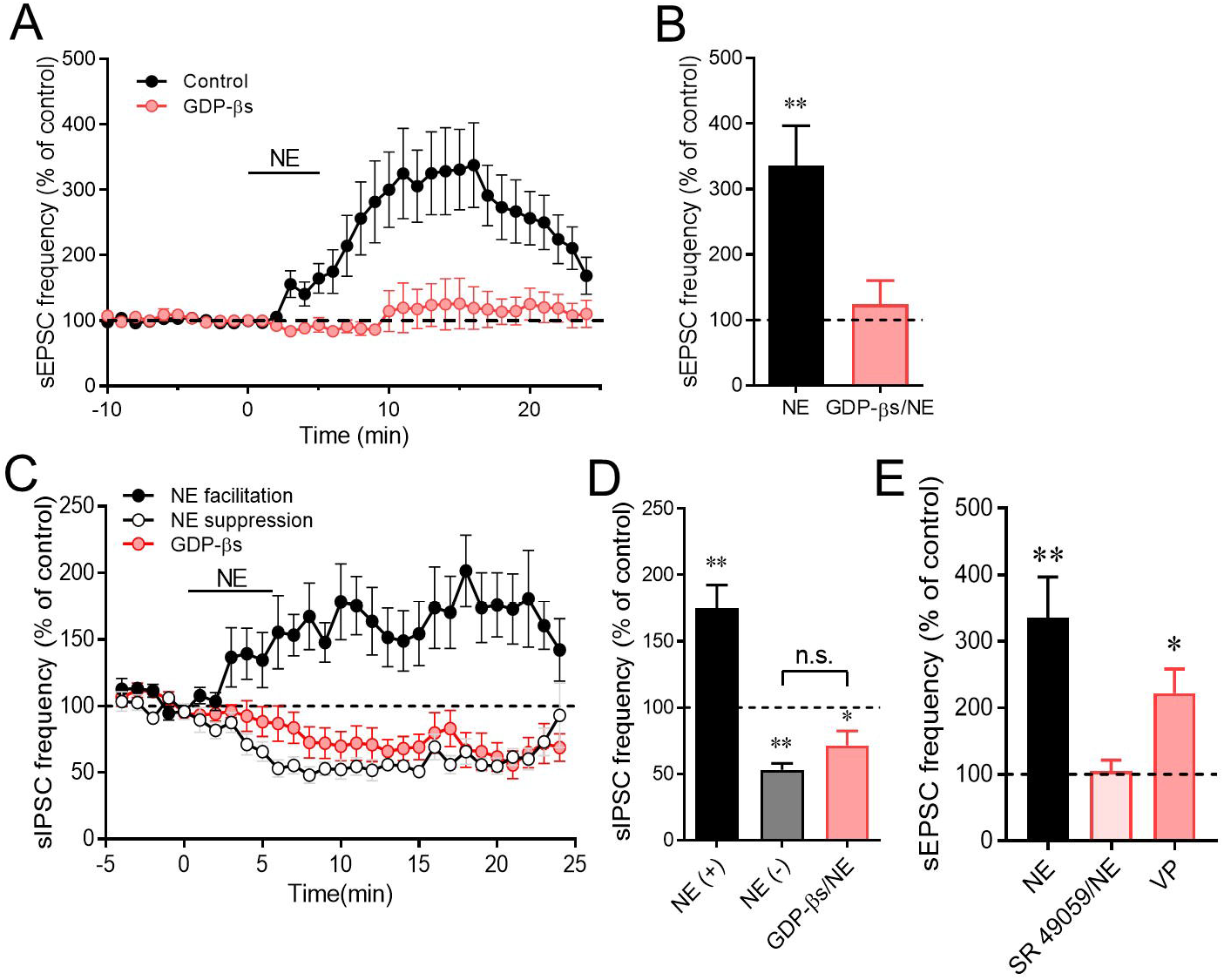
Norepinephrine regulation of synaptic inputs to CRF neurons requires dendritic release of vasopressin. A. The NE-induced increase in sEPSC frequency is blocked by intracellular application of a G-protein blocker, GDP-βs, in the CRF neurons. B. Average sEPSC frequency changes in response to NE in control CRF neurons (NE) and in CRF neurons in which G-protein activity has been blocked with GDP-βs. C. Time plots of the NE-induced increase in sIPSC frequency, which is blocked by blocking postsynaptic G protein activity, and decrease in sIPSC frequency, which is resistant to postsynaptic G-protein blockade. D. Average sIPSC frequency facilitation (NE(+)) and suppression (NE(-)) to NE without and with GDP-βs in the patch pipettes. The facilitation of sIPSCs by NE was blocked, but the suppression of sIPSCs was maintained during postsynaptic G protein inhibition. E. The vasopressin V1a receptor antagonist SR 49059 blocked the NE-induced increase in sEPSC frequency and puff application of vasopressin (VP) elicited a similar increase in sEPSC frequency.

### The norepinephrine-induced facilitation of excitatory and inhibitory synaptic inputs to CRF neurons is mediated by dendritic vasopressin release

The activation of presynaptic glutamate and GABA neurons by a postsynaptic α_1_ adrenoreceptor-dependent mechanism implicates the dendritic release of an excitatory retrograde messenger. We tested for the involvement of nitric oxide (NO) and CRF, but the NE effect was not dependent on either signal (Supplementary Fig. 3). Vasopressin is expressed in CRF neurons and released from CRF neuron axon terminals in the median eminence in response to different stressors and central α_1_-adrenoreceptor activation (Whitnall et al., 1993; de Goeij et al., 1991). We showed recently that vasopressin is released as a retrograde messenger from the dendrites of magnocellular vasopressinergic neurons in the PVN and activates a retrograde signaling mechanism that increases spike-dependent GABAergic inputs to the vasopressin neurons ^37^. Therefore, we tested for the vasopressin dependence of the NE-induced facilitation of sEPSCs. While without effect on basal sEPSCs (p = 0.42), bath application of the vasopressin V1a receptor antagonist SR 49059 completely blocked the increase in sEPSC frequency induced by 100 μM NE (104.5% ± 17.07% of baseline, n = 6, p = 0.80) (Fig. 3E). We then tested for an agonistic effect of vasopressin on sEPSC frequency with pressure application of vasopressin (20 μM) on the surface of the slices close to the recorded CRF neurons (8-20 psi, 30 s). Vasopressin caused a robust increase in the sEPSC frequency in CRF neurons (221.9% ± 36.56%, n = 6, p = 0.021) (Fig. 3E), which was not significantly different from the NE-induced increase in sEPSC frequency. These two experiments suggested that vasopressin is the messenger that is released in response to NE and that activates presynaptic glutamate neurons. However, the release of vasopressin from CRF neuron dendrites has not been described before and is unexpected, so we tested whether the source of vasopressin was from neighboring vasopressin neurons in the PVN. In our previous study, we found that ghrelin elicits the dendritic release of vasopressin from vasopressin neurons in the PVN, resulting in an increase in TTX-sensitive synaptic inputs to the vasopressin neurons ^37^. Bath application of ghrelin (100 nM) had no effect on the sEPSCs recorded in CRF neurons (frequency: 104.5% ± 11.54% of baseline, n = 10, p = 0.71) (Supplementary Fig. 3B), which excluded vasopressin neurons as the source of intra-PVN vasopressin and suggested that the V1a receptor-dependent NE response was mediated by vasopressin release from CRF neurons.

### The norepinephrine facilitation of excitatory and inhibitory inputs to CRF neurons is dependent on astrocyte activity and gliotransmission

The spike dependence of the retrograde facilitation of excitatory and inhibitory synaptic inputs to the CRF neurons suggests that the dendritic messenger acts at a distal presynaptic somatic/dendritic locus to excite presynaptic glutamate and GABA neurons. We tested for a role of astrocytes as intermediates in the relay of the retrograde signal responsible for the NE-induced activation of presynaptic excitatory and inhibitory neurons. Fluorocitric acid (FCA) is preferentially taken up by glia and reversibly blocks glial metabolic function by impairing the Krebs cycle (Paulsen et al., 1987; Swanson and Graham, 1994), which inhibits glial signaling. Preincubation of slices in FCA (100 µM) for 2-4 h prior to recordings significantly blunted the NE-induced increase in sEPSC frequency (144.33% ± 16.23%, n = 13, p = 0.018 compared to baseline, p < 0.01 compared to NE effect in untreated slices) (Fig 4A, B). Preincubation of slices in FCA also blocked the NE-induced increase in sIPSC frequency, resulting in a decrease in sIPSC frequency (74.74% ± 7.78% compared to baseline, n = 10, p = 0.01) (Fig. 4C, D). This suggested that the NE-induced facilitation of both excitatory and inhibitory synaptic inputs to CRF neurons is dependent on astrocyte activity.

**Figure 4.**
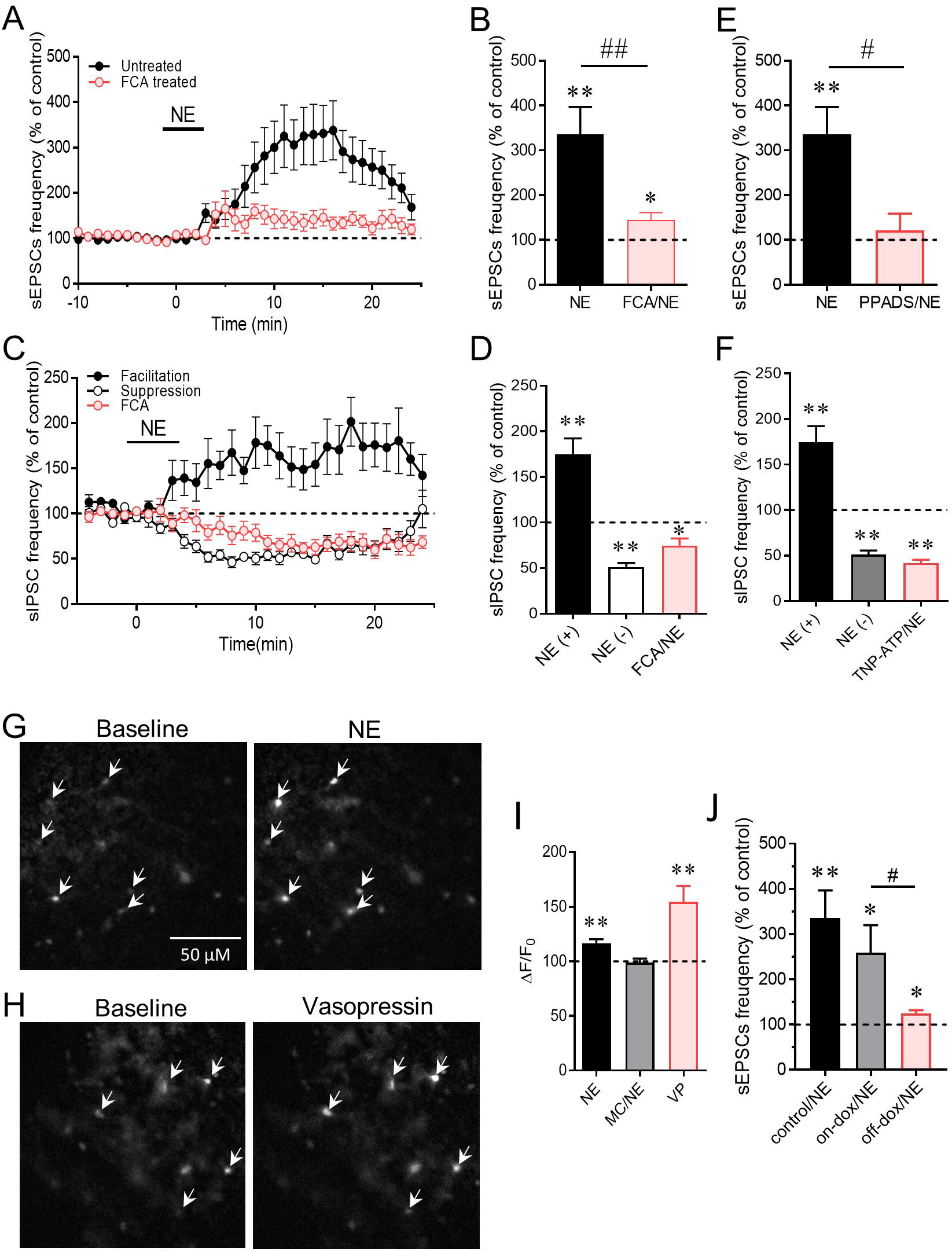
Dependence of the NE effects on astrocytes. A. The NE-induced increase in sEPSC frequency is inhibited by preincubation of slices in the gliotoxin FCA. B. Mean sEPSC frequency changes in NE (100 μM) relative to baseline in control slices and slices incubated in FCA. C. The NE-induced increase in sIPSC frequency is sensitive to the gliotoxin FCA, but not the NE-induced decrease in sIPSC frequency. D. Mean NE-induced increase (NE(+)) and decrease (NE(-)) in sIPSC frequency relative to baseline following FCA treatment. The NE-induced increase in sIPSC frequency (NE(+)), but not the decrease in sIPSC frequency (NE(-)), was blocked by FCA preincubation. E. The NE-induced increase in sEPSC frequency was blocked by the P2 purinergic receptor antagonist PPADS. F. The NE-induced increase in sIPSC frequency was blocked by the P2x purinergic receptor antagonist, TNP-ATP, but the NE-induced decrease in sIPSC frequency was not. G. Fluorescence image showing the glial Ca^2+^ response to NE in the PVN with a glia-specific Rhod-2 AM Ca^2+^ indicator. H. Fluorescence image of the glial Ca^2+^ response to vasopressin. I. Quantification of glial Ca^2+^ responses to NE, which is blocked by the vasopressin receptor blocker Manning Compound (NE/MC), and to vasopressin (VP). J. The NE effect on sEPSCs is maintained in Tet-off dn-SNARE mice maintained on a doxycycline (Dox) diet, partially blocked following 3 weeks of removal of Dox diet, and almost completely lost following 5 weeks off the Dox diet. *, p < 0.05; **, p < 0.01; #, p < 0.05 and ##, p < 0.01 vs. indicated groups.

We next tested whether the NE-induced increase in sEPSC and sIPSC frequencies is dependent on the release of the gliotransmitter ATP by blocking purinergic receptors. Bath application of PPADS (100 µM), a non-selective P2 receptor antagonist, blocked the NE-induced increase in sEPSC frequency (120.8 ± 37.7% of baseline, n = 7, p = 0.42; NE vs. NE/PPADS: p = 0.04) (Fig 4E). In recordings of sIPSCs, bath application of TNPATP (10 μM), a selective P_2_X purinergic receptor antagonist, blocked the NE-induced increase in sIPSC frequency, sparing the NE-induced decrease in sIPSC frequency (41.89% ± 3.57%, n = 6, p < 0.01) (Fig. 4F). The P_2_X antagonist had no effect on the NE-induced decrease in sIPSC frequency (NE vs. TNP-ATP/NE: p = 0.12) (Fig. 4F), which was consistent with the NE-induced suppression of GABA release not being dependent on the retrograde signaling mechanism. Thus, the NE-induced facilitation of excitatory and inhibitory synaptic inputs to the CRF neurons depends on the ATP activation of P_2_X purinergic receptors.

To further test for the glial participation in the NE-induced facilitation of excitatory and inhibitory synaptic inputs, we next conducted calcium imaging experiments using the glia-specific calcium indicator Rhod-2 AM ^40^ to examine whether NE causes a calcium response in glial cells. Slices were incubated in Rhod-2 AM (1-3 μM) for 30-60 minutes to bulk load the calcium fluorophore. Norepinephrine (100 μM, 5 min) induced a significant increase in the relative fluorescence intensity (116.4% ± 3.88%, n = 30, p<0.01), which was blocked with a non-selective oxytocin and vasopressin receptor antagonist, Manning compound (kindly provided by Professor Maurice Manning, University of Toledo) (Fig. 4G, I). We did not use the selective V1a receptor antagonist here because it induced autofluorescence in the brain slices. We next tested whether vasopressin activates a calcium response in astrocytes. Bath application of vasopressin (200 nM) also caused an increase in the relative fluorescence signal (154.7% ± 14.38%, n = 9, p < 0.01) (Fig. 4H, I). These results together suggest that NE stimulates presynaptic glutamate and GABA circuits by activating PVN astrocytes via a vasopressin V1a receptor-dependent mechanism.

The recruitment of glia into the retrograde signaling mechanism triggered by NE and the involvement of ATP as a gliotransmitter in the activation of presynaptic glutamate and GABA neurons suggested that ATP may be released from astrocytes by exocytosis. We tested this hypothesis using a transgenic mouse model (kindly provided by Dr. Phillip Haydon, Tufts University) in which exocytosis is suppressed in astrocytes by the conditional Tet-off expression of a dominant-negative synaptotagmin (dnSNARE) under the control of the glial fibrillary acidic protein promoter. The dnSNARE mice were crossed with the CRF-eGFP mice to allow us to target the CRF neurons for recording. Mice were taken off the doxycycline diet for 5 weeks to induce dnSNARE expression, and mice maintained on the doxycycline diet to suppress dnSNARE expression served as controls. The NE-induced increase in sEPSC frequency was maintained in CRF neurons from CRF-eGFP/dnSNARE mice kept on the doxycycline diet (on-dox/NE: 258.7% ± 60.85% compared to baseline, n = 6, p = 0.048; on-dox/NE vs. control/NE: p = 0.50). In slices from CRF-eGFP/dnSNARE mice taken off the doxycycline diet, the NE-induced increase in sEPSC frequency, although still significant compared to baseline (124.4% ± 7.27%, n = 7, p = 0.015), was significantly suppressed compared to the NE effect in slices from mice maintained on the doxycycline diet (off-dox/NE vs. on-dox/NE: p = 0.037) (Fig. 4J). This suggested that the NE-induced facilitation of glutamate release onto CRF neurons is suppressed by blocking exocytosis in astrocytes.

### Modulation of excitatory and inhibitory synaptic inputs to CRF neurons by endogenous norepinephrine

To determine whether endogenous NE release exerts a similar modulatory effect on excitatory and inhibitory synaptic inputs to PVN CRF neurons, we applied an optogenetic strategy to activate noradrenergic afferent inputs. An AAV9 expressing Cre-dependent channelrhodopsin and mCherry was injected bilaterally into the nucleus of the solitary tract (NTS) of CRF-eGFP, TH-Cre mice. Following confirmation using confocal microscopy of mCherry expression in cell bodies of the NTS and in axon terminal fields in the PVN (Fig. 5A, B), we tested for an excitatory synaptic response in CRF-eGFP neurons to photostimulation of the PVN with blue light (490 nm, 2 min continuous) to activate channelrhodopsin. CRF neurons responded to photostimulation with an increase in sEPSC frequency (355.4% ± 65.8%, n=12, p < 0.01) (Fig. 5C, D), but not amplitude or decay (p = 0.99, 0.89, respectively). The increase in sEPSC frequency was significantly inhibited by the α_1_-adrenoreceptor antagonist prazosin (10 μM) (p = 0.041), but a significant EPSC frequency response to the photostimulation remained (243.4% ± 44.68%, n = 7, p = 0.018) (Fig. 5D). Similarly, the increase in sEPSC frequency to photostimulation was significantly inhibited by blocking postsynaptic G protein activity in the recorded cells with GDP-β-S application (1 mM) via the patch pipette (p = 0.03), but the effect of photostimulation was not completely blocked, as a significant increase in sEPSC frequency remained (252.5% ± 35.15%, n = 6, p < 0.01) (Fig. 5D). The α1 receptor- and G protein-independent response suggests that glutamate may be co-released from the activated noradrenergic afferents.

**Figure 5.**
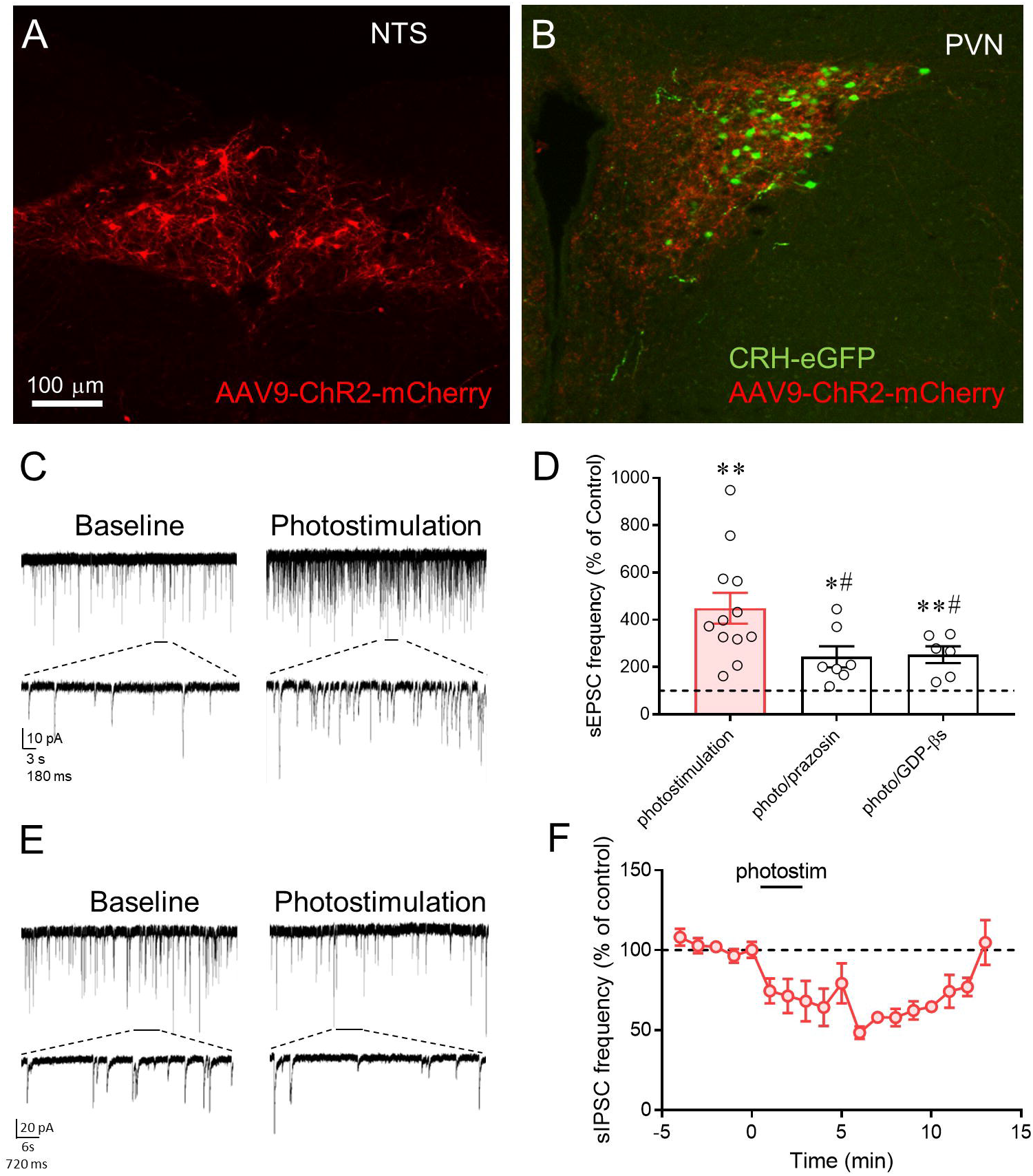
Endogenous norepinephrine modulation of excitatory and inhibitory synaptic inputs to the CRF neurons. A. ChR2-mCherry expression in TH-expressing neurons in the NTS. B. CRF-eGFP expression in neurons and ChR2-mCherry expression in TH-containing axons in the PVN. C. Recording of sEPSCs in a CRF-eGFP neuron in the PVN before (Baseline) and after photostimulation of ChR2 in the presence of GABA_A_ receptor antagonist. D. The sEPSC frequency response to photostimulation before and after blocking α1 receptors with prazosin and postsynaptic G protein activity with intracellular GDP-βs. E. Recording of sIPSCs in a CRF-eGFP neuron before (Baseline) and after photostimulation of ChR2 in the presence of glutamate receptor antagonists. F. Time plot of sIPSC frequency response to photostimulation relative to baseline frequency.

We also tested for the modulation of IPSCs by endogenous NE release in slices from CRF-eGFP, TH-cre mice following virally-mediated channelrhodopsin transduction in the NTS. In the presence of glutamate receptor antagonists, photostimulation failed to elicit a facilitation of sIPSCs, but caused a decrease in sIPSC frequency in all the cells tested (48.45% ± 3.95% of baseline, n = 6, p < 0.01) (Fig. 5E, F).

## Discussion

We found that NE stimulates CRF neurons by a novel dendritic signaling mechanism that activates upstream presynaptic glutamate and GABA neurons via transmission through astrocytes, as shown in our model in Fig. 6. The astrocyte participation in the retrograde signaling significantly expands the spatial domain of the signal to more distal targets than would be reached by dendritic volume transmission alone. This retrograde neuronal-glial signaling mechanism is novel because the signaling is not spatially restricted to the three components of the tripartite synapse, i.e., the presynaptic terminal, the postsynaptic dendrite, and the peri-synaptic astrocyte branch ^41^. This retrograde signaling spans a potentially significant distance to the presynaptic somata/dendrites of upstream glutamate and GABA neurons to drive action potential generation. Indeed, the spatial reach of the retrograde signal could attain even greater distances if the astrocytic calcium signal were transmitted to coupled astrocytes via gap junctions. This needs to be further studied, but could explain why we saw a less robust NE facilitation of inhibitory than excitatory synaptic inputs, since the presynaptic GABA neurons are likely to be located outside the PVN (Boudaba et al., 1996; Herman et al., 2004) (but see also Jiang et al., 2018), while the presynaptic glutamate neurons may be intrinsic to the PVN and thus closer to the CRF neurons (Daftary et al., 1998; Daftary et al., 2000; Wittmann et al., 2005; Hrabovszky et al., 2005; Hrabovszky et al., 2005). Consistent with a difference in the relative distances of presynaptic glutamate and GABA neurons from the CRF neurons, the facilitation of EPSCs was seen in over 90% of recorded CRF neurons, whereas the facilitation of IPSCs occurred in 55% of the neurons, suggesting that the presynaptic GABA neurons are more distal and fewer retain intact axonal projections to the CRF neurons in brain slices. Also, the weaker facilitation of IPSCs than EPSCs by NE (x⎕ ≈ 75% vs. 300%, respectively, at 100 μM) may be due to the lower fidelity of the retrograde signaling to presynaptic GABA neurons because of their remote extra-nuclear location. Signaling to neurons outside the PVN is likely to require transmission through more than one astrocyte, since PVN astrocytes appear to be contained largely within the nucleus. Our findings using optogenetics also support a difference in the locations of the presynaptic glutamate and GABA neurons.

**Figure 6.**
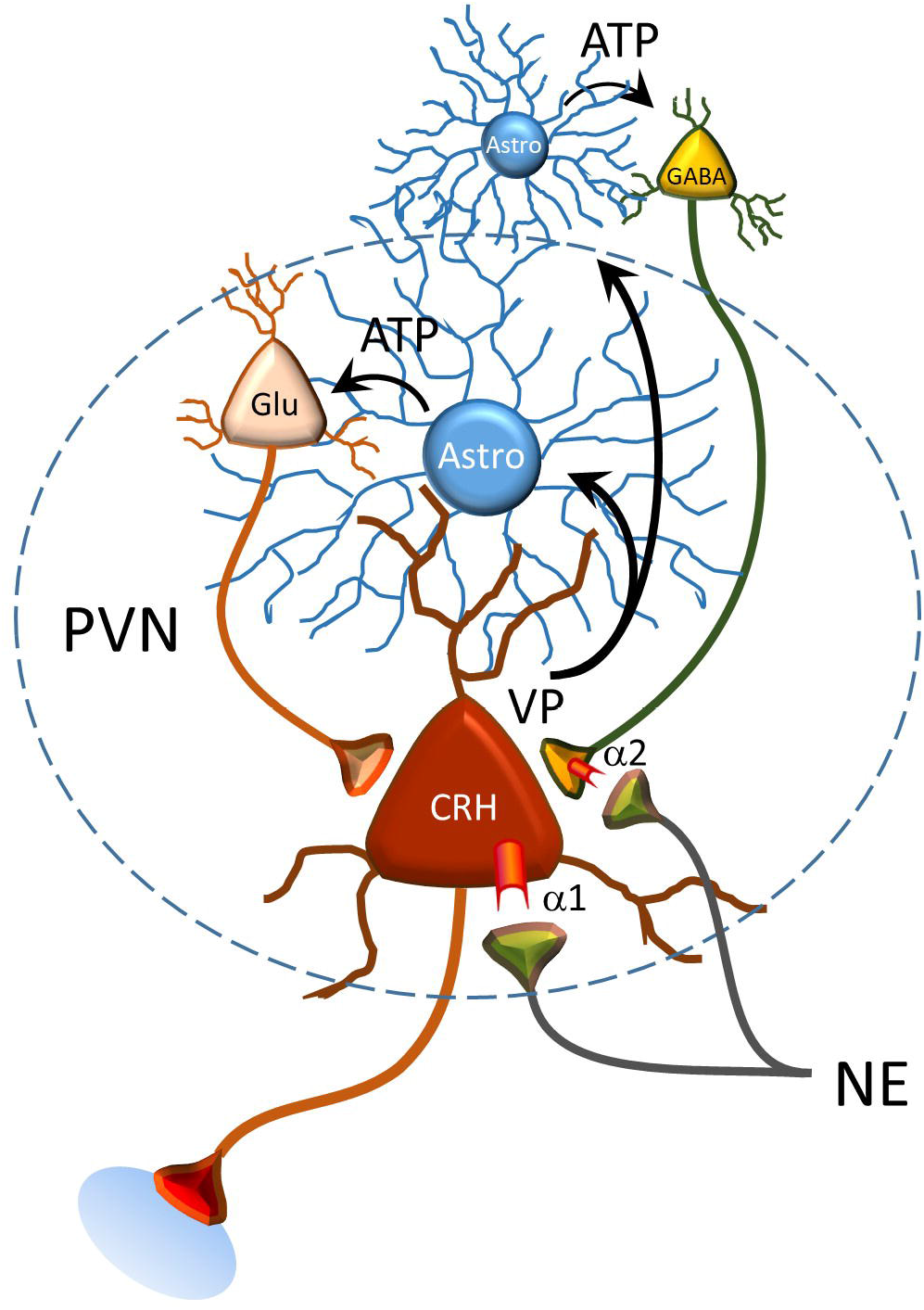
Model of the noradrenergic regulation of CRF neurons. Norepinephrine activates postsynaptic α1 adrenoreceptors on CRF neurons and presynaptic α2 receptors on GABA terminals. α1 receptor activation triggers the dendritic release of vasopressin, which stimulates the astrocytic release of ATP. ATP acts on upstream glutamate and GABA neurons to stimulate spiking, which activates recurrent excitatory and inhibitory inputs to the CRF neurons. Presynaptic α2 receptor activation suppresses the release of GABA. The combined activation of presynaptic glutamate neurons and suppression of GABA release causes excitation of the CRF neurons, which is constrained at higher concentrations of NE by activation of presynaptic GABA neurons.

Photostimulation of noradrenergic axons elicited an α1 receptor-dependent increase in glutamatergic synaptic inputs, but failed to facilitate GABAergic inputs, which is likely due to a higher threshold of activation of the retrograde signaling pathway to the presynaptic GABA neurons because of their extranuclear location and polysynaptic neuronal-glial pathway. The rightward shift in the NE concentration required to activate GABAergic inputs compared to glutamatergic inputs is consistent with a higher threshold to activate the presynaptic GABA neurons. The component of the photostimulated response that was blocked by the α1 antagonist (~50%) (Fig. 5D) suggests that the effective concentration of released NE was between 100 nM and 1 μM, which was subthreshold for the facilitatory effect of NE on GABA release (see Fig. 1).

We found converging lines of evidence for an astrocyte intermediate in the retrograde transmission to presynaptic glutamate and GABA neurons. First, both NE and vasopressin elicited calcium responses in astrocytes. Second, the NE facilitation of synaptic inputs to the CRF neurons was lost following pretreatment of brain slices with the gliotoxin FCA. Third, the NE responses were also lost in CRF neurons from transgenic mice in which exocytosis was conditionally suppressed in astrocytes. Finally, the NE responses were blocked by antagonists of the conventional gliotransmitter, ATP. The sensitivity of the responses to the P2x receptor antagonists and to TTX, and their dependence on astrocytic exocytosis, suggest that astrocytic ATP release drives action potential generation in presynaptic glutamate and GABA neurons by activating purinergic receptors, consistent with our previous findings ^37^. Thus, the evidence for the retrograde transmission to presynaptic glutamate and GABA neurons by way of one or more intercalated astrocytes is compelling.

A surprising discovery of this study was that the retrograde transmitter is vasopressin, which was indicated by the blockade of both the NE-induced synaptic response in CRF neurons and the calcium response in astrocytes by vasopressin receptor antagonists, but not by a CRFR1 antagonist or NO scavenger, and by a vasopressin-evoked synaptic response in the CRF neurons and calcium response in astrocytes. Astrocytes express vasopressin V1a receptors (Hatton et al., 1992; Yamazaki et al., 1997) and we showed previously that vasopressin generates a calcium response in PVN astrocytes ^37^. The source of the vasopressin could have been from neighboring PVN vasopressin neurons, but this was ruled out because ghrelin, which stimulates vasopressin release from vasopressin neuron dendrites ^37^, had no effect on sEPSCs in the CRF neurons.

CRF neurons express vasopressin mRNA and peptide in their somata and axons basally (Whitnall et al., 1987; Lightman and Young, 1988) and increase their vasopressin expression following acute stress, chronic stress (Bartanusz et al., 1993; Baitanusz et al., 1993; Ma et al., 1997), and adrenalectomy ^55^. Vasopressin is co-released with CRF in the median eminence and facilitates adrenocorticotrophic hormone secretion from the pituitary ^56^. Our findings support previous studies showing that α1-adrenoceptor activation stimulates a vasopressin-containing subpopulation of CRF neurosecretory cells (Cummings and Seybold, 1988; Whitnall et al., 1993), and indicate that vasopressin is also released from CRF neuron dendrites.

Norepinephrine induced an α2 receptor-dependent suppression of sIPSCs in 45% of recorded neurons, which increased to 100% when the α1 receptor-mediated facilitation was blocked with TTX. Thus, the α2 receptor suppression of GABA release was masked in a subset of CRF neurons (55%) at high NE concentration by α1 receptor-dependent stimulation of presynaptic GABA neurons with intact afferent axons. The two GABA responses to NE had differing concentration-dependencies, with the GABA suppression effective at lower concentration (1 μM) and the facilitation emergent at higher concentrations (>10 μM). The threshold concentration for the NE facilitation of excitatory inputs was lower than both, near 100 nM. The results of optogenetic stimulation are consistent with the lower threshold of NE activation of local excitatory circuits compared to inhibitory circuits, since photostimulation of the noradrenergic axons elicited a facilitation of excitatory, but not inhibitory, synaptic inputs to the CRF neurons. The combined α1 facilitation of excitatory inputs and α2 suppression of inhibitory inputs by endogenous NE would lead to a robust activation of the CRF neurons and the HPA axis. The higher threshold for the facilitation of GABA inputs may result in the recruitment of inhibitory circuits with more robust activation of the ascending noradrenergic afferents, possibly to prevent excessive activation of the HPA axis.

Our findings reconcile discrepancies between previous neuroanatomical evidence of noradrenergic synapses directly on and α1 adrenoreceptor expression in CRF neurons (Liposits et al., 1986; Cummings and Seybold, 1988; Flak et al., 2009; Cunningham and Sawchenko, 1988; Sawchenko and Swanson, 1982; Day et al., 1999; Cunningham et al., 1990; Füzesi et al., 2007) and electrophysiological data showing an indirect NE regulation of PVN parvocellular neurons via activation of local glutamate and GABA circuits (Daftary et al., 2000; Han et al., 2002). What functional utility is served by a complex indirect regulation of CRF neurons via retrograde activation of local synaptic circuits? The local circuits may represent a common network by which both ascending physiological inputs and descending psychological inputs regulate the HPA axis. While the ascending physiological inputs activate the circuits via retrograde signaling, the descending limbic inputs could activate the same circuits directly. The complex heterotypic nature of the local neuronal-glial organization provides multiple levels of modulation to regulate the circuit.

Retrograde dendritic signaling by transmission via astrocytes and astrocytic networks represents a powerful mechanism for the control by postsynaptic neurons of presynaptic neuronal ensembles, and significantly extends the domain of influence of the postsynaptic neuron over upstream afferent circuits. Whether non-neuroendocrine neurons are also capable of dendritic volume transmission dependent on astrocyte activation is not known. Neuronal-glial signaling has been described in hippocampal neurons via excitatory endocannabinoid actions on astrocytes (Navarrete and Araque, 2008; Navarrete and Araque, 2010) and in dorsal root ganglion neurons via ATP activation of satellite cells “sandwiched” between DRG somata ^61^, but whether these forms of trans-cellular signaling can recruit distant presynaptic neurons remains to be determined.

## Acknowledgments

This work was supported by NIH R01 MH069879 and the Catherine and Hunter Pierson Chair in Neuroscience. We thank Dr. Laura Harrison for her help with the animal husbandry and molecular analyses, and Dr. Diankun Yu for his helpful discussion of the paper. We thank Prof. Maurice Manning for his donation of vasopressin receptor antagonist and Dr. Paul Sawchenko for his donation of CRF antibody.

Author Contributions
C.C. and Z.J. contributed equally to the experiments, data analysis, and to writing the manuscript; X.F. contributed to the experiments and edition of the manuscript; J.G.T. contributed to the design and interpretation of experiments and to writing the manuscript.

